# Glucocorticoid receptor dysregulation underlies 5-HT_2A_ receptor-dependent synaptic and behavioral deficits in a mouse neurodevelopmental disorder model

**DOI:** 10.1101/2022.01.07.475437

**Authors:** Justin M. Saunders, Carolina Muguruza, Salvador Sierra, José L. Moreno, Luis F. Callado, J. Javier Meana, Patrick M. Beardsley, Javier González-Maeso

**Affiliations:** Department of Physiology and Biophysics, Virginia Commonwealth University School of Medicine, Richmond, VA 23298, USA; Department of Pharmacology and Toxicology, Virginia Commonwealth University School of Medicine, Richmond, VA 23298, USA; Center for Biomarker Research and Precision Medicine, Virginia Commonwealth University School of Pharmacy, Richmond, VA 23298, USA; Department of Pharmacology, University of the Basque Country UPV/EHU, CIBERSAM, Biocruces Bizkaia Health Research Institute, E-48940 Leioa, Bizkaia, Spain; Department of Neurology, University of Michigan, Ann Arbor, MI 48109

## Abstract

Prenatal environmental insults increase the risk of neurodevelopmental psychiatric conditions in the offspring. Structural modifications of dendritic spines are central to brain development and plasticity. Using maternal immune activation (MIA) as a rodent model of prenatal environmental insult, previous results have reported dendritic structural deficits in the frontal cortex. However, very little is known about the molecular mechanism underlying MIA-induced synaptic structural alterations in the offspring. Using prenatal (E12.5) injection with poly-(I:C) as a mouse MIA model, we show that upregulation of the serotonin 5-HT_2A_ receptor (5-HT_2A_R) is at least in part responsible for some of the effects of prenatal insults on frontal cortex dendritic spine structure and sensorimotor gating processes. Mechanistically, we report that this upregulation of frontal cortex 5-HT_2A_R expression is associated with MIA-induced reduction of nuclear translocation of the glucocorticoid receptor (GR) and, consequently, a decrease in the enrichment of GR at the *5-HT_2A_R* promoter. The translational significance of these preclinical findings is supported by data in postmortem human brain samples suggesting dysregulated nuclear GR translocation in frontal cortex of schizophrenia subjects. Repeated (twice a day for 4 days) corticosterone administration augmented frontal cortex *5-HT_2A_R* expression and reduced GR binding to the *5-HT_2A_R* promoter. However, virally (AAV)-mediated augmentation of GR function reduced frontal cortex *5-HT_2A_R* expression and improved sensorimotor gating processes via 5-HT_2A_R. Together, these data support a negative regulatory relationship between GR signaling and *5-HT_2A_R* expression in mouse frontal cortex that may carry implications for the pathophysiology underlying *5-HT_2A_R* dysregulation in neurodevelopmental psychiatric disorders.

## INTRODUCTION

Neurodevelopmental psychiatric disorders, such as schizophrenia and autism, are severe and usually cause life-long disability [1, 2]. Epidemiological studies have indicated that maternal infection during pregnancy with a wide variety of agents, including viruses (influenza [3–6] and rubella [7]), bacteria (bronchopneumonia) [8, 9] and protozoa (*Toxoplasma gondii*) [10] increase the risk of neurodevelopmental psychiatric conditions in the adult offspring. Maternal adverse life events that occur during pregnancy, such as war [11, 12], famine [13], and death or illness in a first-degree relative [14], have also been associated with an elevated risk of schizophrenia and autism. The current consensus is that the effects of prenatal insults are likely explained by their capacity to affect the maternal immune system [15, 16]. Accordingly, converging lines of evidence from humans [5] as well as rodent [17] and more recently non-human primate [18] animal models suggest a causal relationship between maternal immune activation (MIA) during pregnancy and neuropathological/behavioral abnormalities consistent with a range of neurodevelopmental psychiatric disorders. Previous results based on microarray and RNA-seq assays have shown changes in gene expression levels that accompany behavioral alterations in rodent MIA models [19–21]. However, there is not presently a clear study assessing the translatability of any of these MIA-affected genes in offspring deficits relevant to neurodevelopmental psychiatric conditions.

The serotonin 5-HT_2A_ receptor (5-HT_2A_R) is a class A G protein-coupled receptor (GPCR) involved in processes related to cognition, perception and mood [22, 23]. Our previous data suggested up-regulation of 5-HT_2A_R in postmortem frontal cortex samples of untreated schizophrenia subjects as compared to individually matched controls [24–27]. Interestingly, we also reported a similar phenotype – *i.e*., upregulation of frontal cortex 5-HT_2A_R density – using three independent mouse MIA models that included maternal infection with a mouse-adapted influenza virus [28, 29], maternal variable stress, and maternal immune activation with poly-(I:C) [30]. This has been followed by numerous reports revealing upregulation of cortical 5-HT_2A_R in rodent models of environmental insults, such as maternal immune activation with poly-(I:C) [31] or lipopolysaccharide (LPS) [32], maternal stress [33, 34], and LPS administration in adult mice [35], which further strengthens the validity of MIA models to replicate biochemical alterations observed in postmortem schizophrenia frontal cortex samples. It remains unknown, however, whether this 5-HT_2A_R upregulation is involved in the negative effects on disorder-related phenotypes such as dendritic structural plasticity and sensory processes. Furthermore, the upstream signaling mechanisms leading to 5-HT_2A_R upregulation have not been resolved.

Dendritic spines are a fundamental component of synaptic structural plasticity with important roles in cognitive and sensory processes [36]. Neuroanatomical studies from postmortem brains of subjects with neurodevelopmental disorders demonstrate altered density and morphology of dendritic spines [37, 38] – particularly in the frontal cortex; a brain region involved in processes related to perception and cognition [39]. Our data suggest that MIA-induced up-regulation of frontal cortex 5-HT_2A_R occurs through a mechanism that requires dysregulation in nuclear localization and consequently binding of the glucocorticoid receptor (GR) to the promoter region of the *5-HT_2A_R* gene. Additionally, we propose that MIA-induced up-regulation of frontal cortex 5-HT_2A_R is at least in part responsible for some of the effects of prenatal insults on frontal cortex dendritic spine structure and sensorimotor gating deficits in the offspring. Together, these findings provide useful information that may serve to design new tools to reduce synaptic structural plasticity deficits and alterations in the regulation of sensory information observed in subjects with neurodevelopmental psychiatric conditions linked to prenatal insults.

## METHODS

### Materials, Drug Administration, and Mouse Brain and Blood Samples

Poly-(I:C) (polyinosinic–polycytidylic acid potassium salt; Millipore Sigma P9582) was dissolved in saline (0.9% NaCl) and, after a brief centrifugation (800 *×*g), the resulting solution (4 mg/ml) was aliquoted and stored at -20 °C until the day of the experiment.

Corticosterone (Millipore Sigma #46148) was diluted in DMSO (100 mg/ml) and aliquots were stored at -20 °C until the day of the experiment. Adult male mice were injected (s.c.) with corticosterone (50 mg/kg) dissolved in a final solution of DMSO:ethanol (1:3). For time-course assays, all mice (excluding the no-injection group) were injected (s.c.) twice a day over eight days with corticosterone (50 mg/kg), or vehicle, and sacrificed 8.5-13 h after the last injection. For the short-term experiments, adult male mice received (s.c.) corticosterone (50 mg/kg), or vehicle, twice a day for four days. Mice were either sacrificed or tested for behavior 8.5-13.5 h after the last injection.

For brain samples, mice were sacrificed for analysis by cervical dislocation, and bilateral frontal cortex (bregma 1.40 to 1.90 mm) was dissected and either frozen at -80 °C or immediately processed for RNA extraction, chromatin immunoprecipitation and/or biochemical assays. The coordinates were taken according to a published atlas of the C57BL/6 and 129Sv mouse brains [40].

For serum samples, following decapitation, trunk blood was collected and allowed to clot at room temperature for at least 30 min. The blood was then centrifuged (1000 *×*g) at 4 °C for 10 min after which the serum supernatant was collected and stored at -80 °C until the day of the assays.

All other chemicals were obtained from standard sources.

### Animals

Experiments were performed on adult (10 – 22 week-old) male C57BL6/N, C57BL6/J and/or 129S6/SvEv mice (details below), unless otherwise indicated. Animals were housed on a 12 h light/dark cycle at 23 °C with food and water ad libitum, except during behavioral testing which took place during the light cycle. All procedures were conducted in accordance with the National Institutes of Health (NIH) guidelines and were approved by the Virginia Commonwealth University Animal Care and Use Committee. Behavioral testing took place between 9:00 a.m. and 6:00 p.m.

5-HT_2A_R (*Htr2a*) knockout (*5-HT_2A_R^-/-^*) mice on a 129S6/SvEv background have been previously described [41]. For experiments involving *5-HT_2A_R^-/-^* mice, wild-type (*5-HT_2A_R^+/+^*) littermates on a 129S6/SvEv background were used as controls. All subjects were offspring of heterozygote breeding.

### Immune Activation with poly-(I:C)

To test the effect of poly-(I:C) administration on female immune response, naïve adult animals (C57BL6/N from Charles River Laboratories, and 129S6/SvEv generated via in-house breeding) were injected (i.p.) with poly-(I:C) (20 mg/kg), or vehicle, and observed for sickness behavior, such as lethargy, ptosis, and hunched posture. After 2.5 h, mice were sacrificed by either decapitation and exsanguination, or by cervical dislocation for collection of frontal cortex samples.

To test the effect of maternal immune activation (MIA) during pregnancy, experiments were carried out with either timed pregnant C57BL6/N mice ordered from Charles River Laboratories, or C57BL6/N females in-house bred with C57BL6/N males from Charles River Laboratories. For MIA experiments in *5-HT_2A_R^-/-^* mice and *5-HT_2A_R^+/+^* controls, 129S6/SvEv female *5-HT_2A_R^+/-^* (heterozygote) mice (Taconic background) were bred in-house; females were monitored for mating plug and pregnancy was evaluated using both weight gain and visual appearance of the flanks. On day E12.5 of pregnancy, mice were injected (i.p.) with poly-(I:C) (20 mg/kg), or vehicle, and then monitored for sickness behavior. This timing was chosen because the first and second trimesters have been shown to be a critical period for increased risk of neurodevelopmental conditions including schizophrenia and autism following influenza virus infection in humans [8], and MIA at E12.5 has been demonstrated to produce MIA-induced phenotypes in offspring [17]. To reduce litter loss, MIA or mock mice were pair-housed with another non-pregnant female of the same strain and provided with enrichment Bio-Huts (Bio-Serv) as shelter in the cage. Offspring were weaned between 3-4 weeks of age, heterozygous *5-HT_2A_R^+/-^* animals were sacrificed, and homozygous *5-HT_2A_R^+/+^* and *5-HT_2A_R^-/-^* littermates were evaluated in biochemical, dendritic spine structure and behavioral assays as adults.

### Prepulse Inhibition of Startle

Prepulse inhibition of startle (PPI) experiments were conducted following a previously described paradigm [42].

### Plasmid Construction

The AAV-CaMKII*α*::eYFP construct has been described previously [43]. The pCMV-GR11 construct was obtained from Addgene (# 89105). To generate the AAV-CaMKII*α*::ΔGR-p2A-eYFP construct, a PCR fragment was obtained from mouse frontal cortex cDNA as a template and the primers mΔGR-BamHI/S and mΔGR-Ascl/A (mΔGR Cloning Fwd and Rev, respectively – Table S1). The resulting PCR product was digested with BamHI and AscI and subcloned into the same restriction sites of AAV-CaMKII*α*-c-Myc-5-HT_2A_R-p2A-eYFP. All PCR construct amplifications were performed with *PfuUltra* high fidelity DNA polymerase (Agilent) in a Mastercycler EP Gradient Auto thermal cycler (Eppendorf). All the constructs were confirmed by DNA sequencing.

### Virally mediated gene transfer

The construct vectors CaMKII*α*::eYFP and CaMKII*α*::ΔGR-p2A-eYFP were packaged into adeno-associated (AAV) serotype 8 vector particles, all produced by the University of North Carolina at Chapel Hill Vector Core. The AAV-eYFP and AAV-ΔGR-eYFP construct vectors were injected into the frontal cortex of adult mice by stereotaxic surgery according to standard methods [43, 44]. Mice were anesthetized with isoflurane (2% initial dose, 1-2% maintenance) during the surgery.

### Dendritic spine analysis

Frontal cortex dendritic spine structural assays were performed as previously reported [43, 44].

### Immunohistochemistry

Experiments in Neuro-2a cells (ATCC: CCL-131) and mouse frontal cortex were performed as previously reported with minor modifications [43]. Briefly, GR immunoreactivity was assayed by using an anti-GR antibody (Santa Cruz sc-393232; 1:50) and Alexa 568 anti-mouse secondary antibody (Invitrogen A-11004; 1:1000) or Alexa-488 anti-mouse secondary antibody (Invitrogen A-11004; 1:2000).

### Fluorescent in situ hybridization

Using a subcloned and sequenced PCR fragment (162 bp – Table S1) of the mouse *5-HT_2A_R* cDNA (Zero Blunt TOPO PCR Cloning kit, Invitrogen), high specific-activity RNA probes were produced from a linearized plasmid (HindIII) and labeled using the Fluorescein RNA Labeling Mix (Roche), following manufacturer’s instructions.

### ELISA assays

Enzyme-linked immunosorbent assays were performed using the mouse IL-6 ELISA-MAX kit (Biolegend), using manufacturer’s instructions.

### Quantitative real-time PCR

Quantitative real-time PCR (qRT-PCR) assays were carried out as previously described [45]. Primer pairs sequences are listed in Table S1 (see also [43, 46–51]).

### Microbiome Evaluation

Adult female mice (C57BL6/N from Charles River Laboratories, C57BL6/N from Taconic, C57BL6/J from Jackson Laboratories, and 129S6/SvEv generated via in-house breeding) were sacrificed and cecum samples were collected based on previously described protocols [28]. DNA was isolated using a PowerFecal DNA Isolation kit (MO BIO Laboratories). For primer pair sequences, see Table S1.

### Cellular fractionation

The cytoplasmic and nuclear fractions from Neuro-2a cells, mouse frontal cortex and postmortem human frontal cortex samples were prepared as previously reported [43].

### Immunoblot assays

Western blot experiments were performed as previously reported, with minor modifications [43]. In Neuro-2a cells and mouse and postmortem human frontal cortex samples, the following primary antibodies were used: mouse anti-GR (Santa Cruz sc-393232, 1:100), mouse anti-*α*-tubulin (Abcam ab7291, 1:3000), rabbit anti-*β*-actin (Abcam ab8227, 1:3000-1:20000).

### Chromatin immunoprecipitation assay

Chromatin immunoprecipitation (ChIP) experiments were performed as previously reported with minor modifications [43, 45], using a MAGnify Chromatin Immunoprecipitation System (Invitrogen).

### Postmortem human brain samples

Human brains were obtained at autopsies performed in the Basque Institute of Legal Medicine, Bilbao, Spain. The study was developed in compliance with policies of research and ethical review boards for postmortem brain studies (Basque Institute of Legal Medicine, Spain). Retrospective searches were conducted for previous medical diagnosis and treatment using examiners’ information and records of hospitals and mental health centers. After searching antemortem information, 32 subjects who had met criteria of schizophrenia according to the Diagnostic and Statistical Manual of Mental Disorders (DSM-IV and/or DSM-IV TR) were selected [52]. Toxicological screening for antipsychotics, other drugs and ethanol was performed in blood, urine, liver and gastric contents samples. The toxicological assays were performed at the National Institute of Toxicology, Madrid, Spain, using a variety of standard procedures including radioimmunoassay, enzymatic immunoassay, high-performance liquid chromatography and gas chromatography–mass spectrometry. The schizophrenia subjects were divided into 16 antipsychotic-free (AP-free) and 16 antipsychotic-treated (AP-treated) subjects according to the presence or absence of antipsychotics in blood at the time of death. Controls for the present study were chosen among the collected brains on the basis, whenever possible, of the following cumulative criteria: (i) negative medical information on the presence of neuropsychiatric disorders or drug abuse and (ii) appropriate sex, age, postmortem delay (time between death and autopsy, PMD) and freezing storage time to match each subject in the schizophrenia group. Causes of death among the schizophrenia patients included suicide (n=19), accidents (n=2) and natural deaths (n=11). Controls’ causes of death included accidents (n=20) and natural deaths (n=12). Specimens of prefrontal cortex (Brodmann’s area 9) were dissected at autopsy (0.5 – 1.0 g tissue) on an ice-cooled surface and immediately stored at −80°C until use. Tissue pH values were within a relatively narrow range (control subjects: 6.48 ± 0.08; schizophrenia subjects: 6.28 ± 0.05). The definitive pairs of AP-free schizophrenia subjects and respective matched controls are shown in Table S2 and the definitive pairs of AP-treated schizophrenia subjects and respective matched controls are shown in Table S3. Pairs of schizophrenia patients and matched controls were processed simultaneously and under the same experimental conditions.

### Statistical analysis

Animals were randomly allocated into the different experimental groups. Statistical significance was assessed by Student’s *t* test, and one-way or two-way (or multi-way) ANOVA, depending upon the number of experimental conditions and independent variables. Following a significant ANOVA, specific comparisons were made using the Bonferroni’s post-hoc test. Correlation analysis was conducted using the Pearson’s *r*. All values in the figures represent mean ± S.E.M. All statistical analyses were performed with GraphPad Prism software version 9, and comparisons were considered statistically significant if p < 0.05.

## RESULTS

Poly-(I:C) is a synthetic analogue of double-stranded (ds)RNA, a molecular pattern associated with viral infections, that is widely used to provoke an immune response in rodent models, including administration of the viral mimetic to activate the immune system of pregnant mice [17]. Recent studies suggest that the immunogenicity and alterations observed in the offspring following administration of poly-(I:C) to pregnant mice may vary in intensity depending on both source and lot of the dsRNA synthetic analogue [53, 54]. To test the effect of poly-(I:C) on the immune response within our experimental system, we tested immunoreactivity and expression of IL-6, a key proinflammatory mediator induced by poly-(I:C) administration [55], in serum and frontal cortex samples of non-pregnant adult 129S6/SvEv (in-house) mice and C57BL6/N mice from Charles River Laboratories (CRL). Mice were administered (i.p.) poly-(I:C) (20 mg/kg) or vehicle, and sacrificed 2.5 h afterwards. Our data show that this model of immune activation leads to an increase in *IL-6* mRNA expression in frontal cortex samples as well as an increase in IL-6 immunoreactivity in serum samples in both 129S6/SvEv (in-house) and C57BL6/N (CRL) animals (Figs. S1A-D). Additionally, poly-(I:C)-treated C57BL6/N (CRL) mice showed reduced body weight 24 h after immune activation (Fig. S1E).

Previous findings have also suggested that maternal gut bacteria composition represents an additional source of variation in neurodevelopmental abnormalities in poly-(I:C)-induced MIA models [46]. Particularly, it has been reported that MIA-induced behavioral alterations were more evident in C57BL6/N mice from Taconic Biosciences (Tac) as compared to C57BL6/J from Jackson laboratories (JAX), and these MIA-induced deficits correlated with the presence of segmented filamentous bacteria (SFB) in the pregnant mother. To evaluate the presence of SFB in female mice, we extracted DNA from cecum contents of 129S6/SvEv (in-house), C57BL6/N (CRL), C57BL6/J (JAX), and C57BL6/N (Tac) animals, and performed qPCR assays targeting the 16S rRNA gene of *SFB*. Consistent with previous findings [46], our data show that C57BL6/J (JAX) animals were *SFB* deficient, whereas presence of *SFB* was more evident in C57BL6/N (CRL) mice as compared to 129SvEv (in-house) and C57BL6/N (Tac) females (Figs. S2A,B). Evaluating the total abundance of the bacterial *16S rRNA* gene using universal qPCR primers that detect this gene across the vast majority of bacterial taxa, we found that C57BL6/N (JAX) mice exhibited a significantly greater abundance of *16S rRNA* gene copies in the cecum as compared to mice from any of the other three sources (Fig. S2C).

Our previous work suggested up-regulation of both 5-HT_2A_R density by radioligand binding assays and *5-HT_2A_R* mRNA expression by qRT-PCR assays in the frontal cortex of adult mice born to mothers infected with a mouse-adapted influenza virus during pregnancy [28, 29]. Using a fluorescence in situ hybridization approach (FISH), our data here show that frontal cortex tissue sections from adult MIA offspring exhibit an increase in *5-HT_2A_R* mRNA relative to mock C57BL6/N offspring mice (Figs. 1A,B). Controls to validate the specificity and selectivity of FISH assay included decreased fluorescent signal upon dilution of the mRNA probe (Figs. S3A,B), as well as absence of fluorescent signal in frontal cortex tissue sections exposed to a sense strand *5-HT_2A_R* mRNA probe (Figs. S3C).

**Fig. 1.**
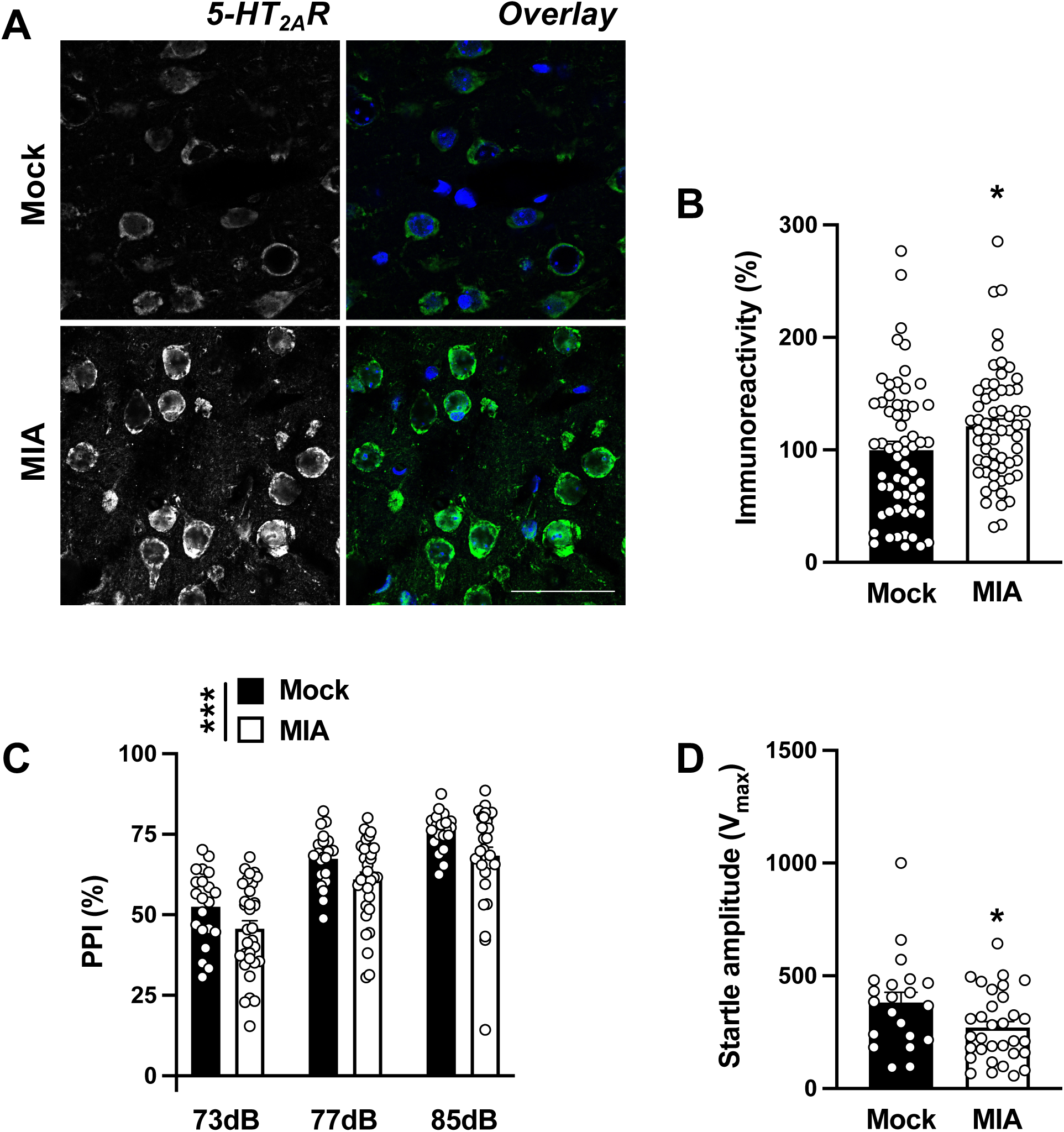
MIA increases frontal cortex *5-HT_2A_R* mRNA expression and disrupts sensorimotor gating in the adult offspring. **A-B,** Increased expression of *5-HT_2A_R* mRNA in the frontal cortex of adult MIA mice. Representative FISH images with the antisense *5-HT_2A_R* probe (**A**), and quantification of fluorescence intensity (n = 61 cells from 4 male C57BL6/N CRL mice/group; 2-3 litters/group) (**B**). **C-D,** Effect of MIA on %PPI (**D**) and startle amplitude (**D**) in the adult offspring (n = 21-32 male and female C57BL6/N CRL mice/group; 3-4 litters/group). Data show mean ± S.E.M. *p < 0.05, **p < 0.01. Unpaired two-tailed student’s *t*-test (**B**: t_120_ = 2.12, p < 0.05; **D**: t_51_ = 2.25, p < 0.05). One-way ANOVA followed by Bonferroni’s post-hoc test (**C:** MIA vs mock F[1, 153] = 12.52, p < 0.001; prepulse intensity F[2, 153] = 46.90, p < 0.001). Nuclei were stained in blue with Hoechst (**A**). Scale bar, 200 µm (**A**).

Prepulse inhibition of the startle is a conserved sensorimotor gating behavioral model that is affected in individuals with certain psychiatric conditions [56]. Similar to previous findings [57], we observed PPI deficits in MIA adult offspring relative to C57BL6/N controls (Fig. 1C). In addition, adult MIA mice exhibited a decrease in startle amplitude during the initial 5 trials of the PPI paradigm (Figs. 1D). Together, these data corroborate that both 129S6/SvEv (in-house) and C57BL6/N (CRL) mice exhibit MIA effects in offspring and therefore serve as a preclinical model to evaluate the effects of MIA on neurodevelopmental processes in the offspring.

Previous findings suggest that MIA affects offspring’s frontal cortex dendritic spine structure [58–60]. To determine whether alterations in frontal cortex 5-HT_2A_R-dependent signaling are involved in the effects of MIA on offspring’s frontal cortex dendritic structural plasticity, heterozygous *5-HT_2A_R^+/-^* 129S6/SvEv pregnant mice were injected with poly-(I:C) or vehicle, and cortical dendritic structure assays were tested in adult *5-HT_2A_R^+/+^* or *5-HT_2A_R^-/-^* offspring (Fig. 2A). To target cortical pyramidal neurons, adult MIA and mock mice were stereotaxically injected into the frontal cortex with adeno-associated virus (AAV8) preparations expressing eYFP under the control of the *CaMKIIα* promoter (Fig. 2B). Using a similar viral vector, our previous findings reported a robust over-expression of eYFP in excitatory pyramidal CaMKII*α*-positive, but not inhibitory parvalbumin-positive neurons [43]. In slices prepared approximately 3 weeks after viral injection, when transgene expression is maximal, this approach enabled us to explicitly discriminate single pyramidal cells and their dendritic spines. Notably, MIA offspring mice showed a selective reduction of mature mushroom spines in CaMKII*α*-positive neurons (Figs. 2C,F). There was also a trend toward MIA effect on stubby spines (Figs. 2C,D) and total spine density (Figs. 2C,G), whereas immature thin spines remained unaffected (Figs. 2C,E). This effect of MIA required expression of 5-HT_2A_R since the frontal cortex synaptic remodeling alteration was not observed in *5-HT_2A_R^-/-^* littermates (Figs. 2C-G). Similar to previous reports [44], adult mice with selective deletion of the *5-HT_2A_R* gene presented a trend towards reduction in the density of frontal cortex dendritic spines (Figs. 2C-G).

**Fig. 2.**
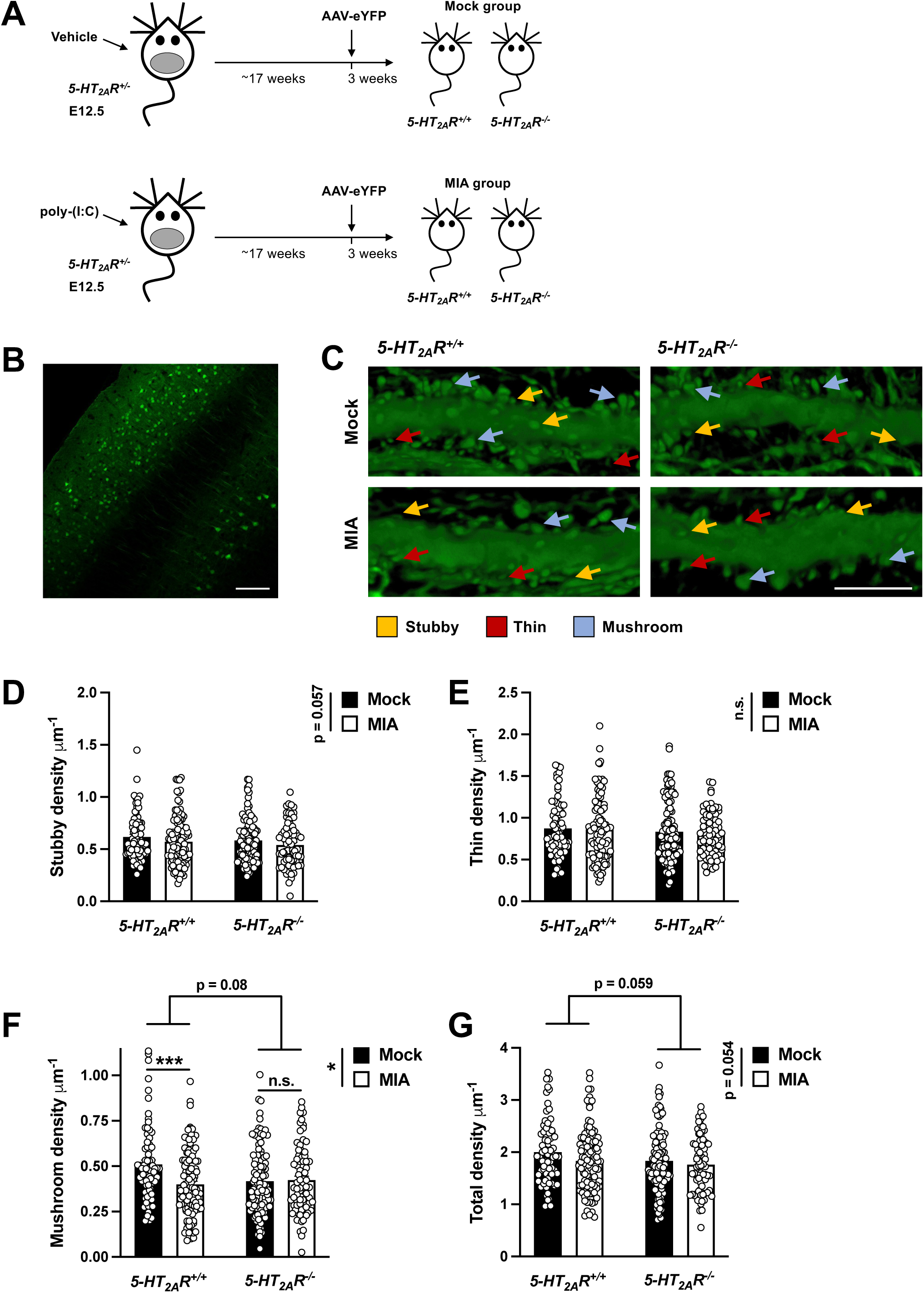
MIA decreases mature synaptic structural elements in mouse frontal cortex via 5-HT_2A_R. **A,** Breeding strategy. Heterozygous *5-HT_2A_R^+/-^* mice were bred to obtain *5-HT_2A_R^+/+^* and *5-HT_2A_R^-/-^* animals from the same mother. Pregnant females (E12.5) received a single injection of poly-(I:C) (20 mg/kg), or vehicle. Adult *5-HT_2A_R^+/+^* and *5-HT_2A_R^-/-^* mice born to either mock or MIA mothers were stereotaxically injected with AAV-eYFP, and sacrificed for analysis 3 weeks after surgery. **B,** eYFP expression by immunohistochemistry. **C,** Representative three-dimensional reconstructions of AAV-injected cortical dendritic segments. **D-G,** Stubby (**D**), thin (**E**), mushroom (**F**) and total (**G**) frontal cortex spine density in adult *5-HT_2A_R^+/+^* and *5-HT_2A_R^-/-^* mice born to either mock or MIA mothers (n = 73-109 neurons from 5-8 male and female 129S6/SvEv in-house mice/group; 3-5 litters/group). Data show mean ± S.E.M. *p < 0.05, ***p < 0.001, n.s., not significant. Two-way ANOVA followed by Bonferroni’s post-hoc test (**D:** stubby MIA vs mock F[1, 363] = 3.64, p = 0.057; stubby genotype F[1, 363] = 1.96, p > 0.05; **E:** thin MIA vs mock F[1, 363] = 0.48, p > 0.05; thin genotype F[1, 363] = 1.97, p > 0.05; **F:** mushroom MIA vs mock F[6, 363] = 6.58, p < 0.05; mushroom genotype F[1, 363] = 3.01, p = 0.08; **G:** total MIA vs mock F[1, 363] = 3.72, p = 0.054; total genotype F[1, 363] = 3.58, p = 0.059). Scale bars, 100 µm (**B**), 5 µm (**C**).

Glucocorticoid receptors (GR), which belong to an evolutionary conserved nuclear receptor superfamily, are involved in the mechanism by which steroid hormones, such as cortisol in most mammals including humans and corticosterone in laboratory rodents, regulate stress responsiveness as ligand-dependent transcription factors across different species [61]. Previous reports raise the possibility that the GR regulates *5-HT_2A_R* transcription in vitro in different experimental systems [62], but the basic molecular mechanism underlying crosstalk between the glucocorticoid and serotonin receptor systems *in vivo* remains unsolved. Using the consensus sequence for the GR obtained from transcription factor binding datasets [63], we detected a predicted GR binding site at the promoter region of the mouse *5-HT_2A_R* gene (Fig. 3A). Based on these findings, we performed a series of chromatin immunoprecipitation followed by quantitative PCR (ChIP-qPCR) assays with an anti-GR antibody in mouse frontal cortex samples throughout different regions of the *5-HT_2A_R* gene (Fig. 3B). Our ChIP-qPCR data revealed enrichment of GR at the predicted binding site, but not at other locations along the *5-HT_2A_R* gene (Fig. 3C).

**Fig. 3.**
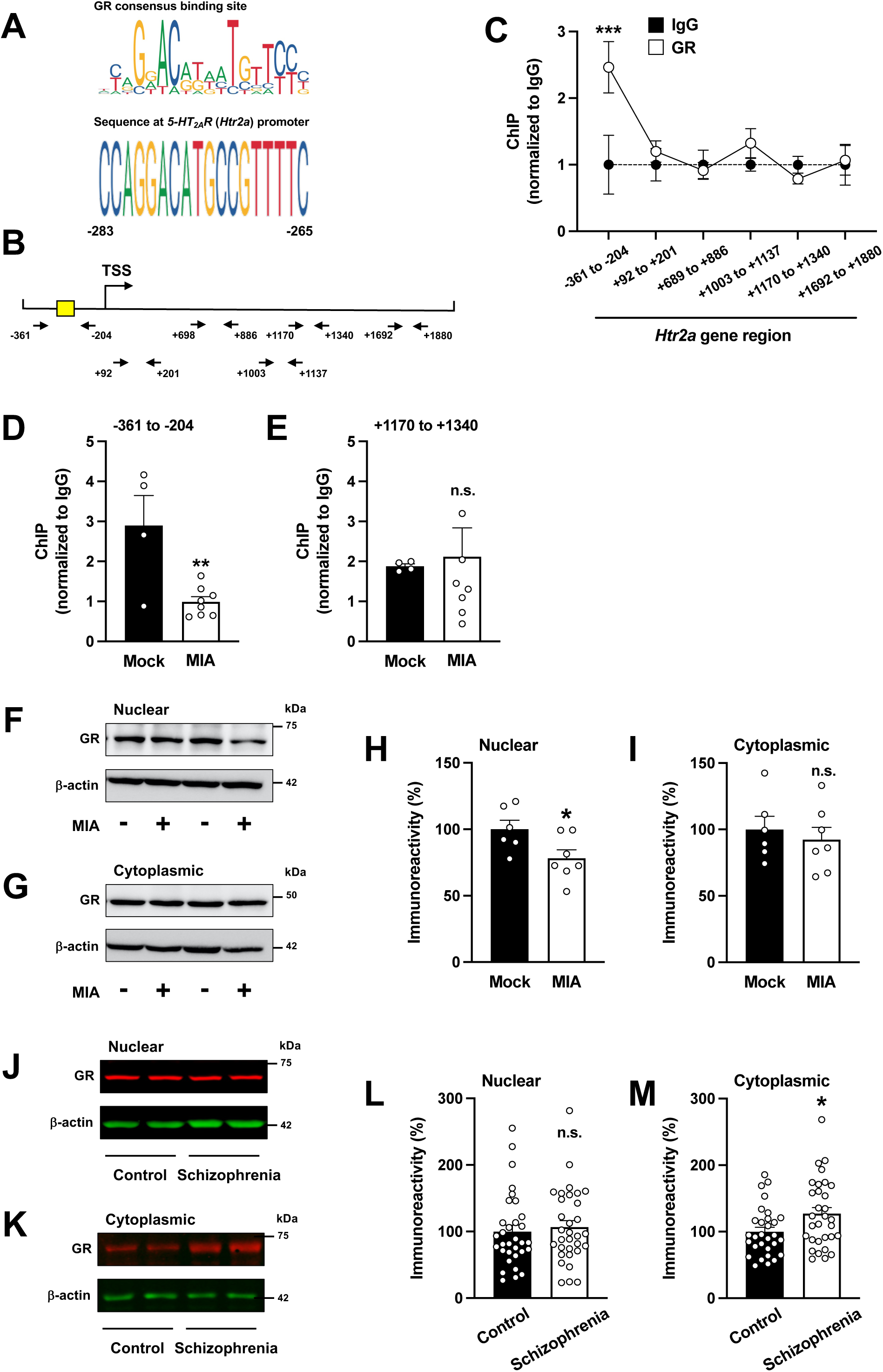
MIA dysregulates localization and binding of GR to the *5-HT_2A_R* promoter in mouse frontal cortex. **A,** Alignment of the vertebrate GR consensus binding site with the promoter of the mouse *5-HT_2A_R* gene. **B,** Map of the *5-HT_2A_R* gene showing position of primers used for qPCR assays. Yellow box indicates location of the predicted GR binding site. TSS, transcriptional start site. **C,** GR binds to the promoter region of the *5-HT_2A_R* gene in mouse frontal cortex (n = 8 male C57BL6/N CRL mice/group). **D-E,** Binding of GR to the region of the *5-HT_2A_R* gene containing a putative GR binding site (**D**), but not within exon 1 (**E**), is decreased in frontal cortex samples of adult MIA mice (n = 4-8 male C57BL6/N CRL mice/group; 3-4 litters/group). **F-I,** Immunoblots showed downregulation of GR protein in nuclear (**H**) but not cytoplasmic (**I**) preparations from the frontal cortex of adult MIA mice as compared to mock animals. Representative immunoblots are shown (**F,G**) (n = 6-7 male C57BL6/N CRL mice/group; 3-4 litters/group). **J-M,** Immunoblots showed upregulation of GR protein in cytoplasmic (**M**) but not nuclear (**L**) preparations from postmortem human brain samples of schizophrenia subjects as compared to controls (for demographic information, see Tables S2 and S3). Representative immunoblots are shown (**J,K**). Data show mean ± S.E.M. *p < 0.05, **p < 0.01, ***p < 0.001 n.s., not significant. Two-way ANOVA followed by Bonferroni’s post-hoc test (**C:** antibody F[1, 73] = 3.83, p = 0.054; gene site F[5, 73] = 2.80, p < 0.05). Unpaired two-tailed student’s *t*-test (**D**: t_10_ = 3.57, p < 0.01; **E:** t_10_ = 0.23, p > 0.05; **H**: t_11_ = 2.34, p < 0.05; **I:** t_11_ = 0.57, p > 0.05; **L:** t_62_ = 0.47, p > 0.05; **M:** t_59_ = 2.35, p < 0.05).

Following this finding in naïve mice, we next sought to determine the effect of MIA on GR enrichment at the *5-HT_2A_R* promoter in mouse frontal cortex. Notably, ChIP-qPCR data reveal that prenatal MIA leads to a significant decrease in the enrichment of GR at the *5-HT_2A_R* promoter (Fig. 3D), but not at a different location within exon 1 (Fig. 3E).

To gain insights into a potential mechanism underlying these findings, and given that activation of the GR pathway can induce GR nuclear translocation, we tested whether prenatal MIA affects subcellular localization of GR in mouse frontal cortex. Consistent with previous studies, GR immunoreactivity was observed in both nuclear and cytoplasmic preparations of frontal cortex samples from control mice (Figs. 3F-I). Importantly, MIA offspring animals showed decreased immunoreactivity against nuclear (Figs. 3F,H), but not cytoplasmic (Figs. 3G,I), GR in the frontal cortex.

To test for a potential dysregulation of GR trafficking in individuals with neurodevelopmental psychiatric conditions, we performed cellular fractionation followed by immunoblot assays with anti-GR antibodies in postmortem frontal cortex samples of schizophrenia subjects and individually matched controls (Tables S2 and S3). Consistent with our findings in mice, we observed a statistically significant difference related to GR subcellular localization in frontal cortex samples from schizophrenia subjects as compared to controls (Figs. 3J-M). This alteration in GR nuclear translocation was also apparent as a trend in both antipsychotic-free and antipsychotic-treated schizophrenics versus their respective control groups (Fig. S4).

Following our data suggesting that MIA produces both up-regulation of *5-HT_2A_R* mRNA expression and decreased binding of GR to the *5-HT_2A_R* promoter in the mouse frontal cortex, we focused our investigation on the effects of direct manipulation of the GR system on 5-HT_2A_R-dependent behavioral phenotypes. Previous investigation has reported that a regimen of corticosterone twice a day for four days produces an increase in head-twitch behavior – a rodent behavioral proxy of human hallucinogenic potential [64] – in response to the psychedelic 5-HT_2A_R agonist DOI in rats [65]. To evaluate and optimize this experimental system in 129S6/SvEv mice, we first administered (s.c.) corticosterone (50 mg/kg) twice a day for up to eight days, and sacrificed 8.5-13 h after the last injection. Control groups included mice injected with vehicle as well as injection-naïve mice. Previous findings have established FKBP5 as an interactor that opposes nuclear translocation of the GR and hence decreases GR-dependent transcriptional activity. Upon glucocorticoid binding, however, FKBP5 is exchanged for FKBP4, a co-chaperone that promotes GR nuclear translocation and transcriptional regulation [66]. Our data suggest that neither *GR* (Fig. S5A) nor *FKBP4* (Fig. S5B) mRNA expression were affected upon repeated corticosterone administration. However, *FKBP5* mRNA (Fig. S5C) showed upregulation in the frontal cortex of mice after either 1, 2, 4, 6 or 8 days of repeated corticosterone administration, as compared to either the vehicle; the injection-naïve group was not significantly different from vehicle. These findings suggest that repeated corticosterone administration activates and elicits negative feedback within the GR pathway in the mouse frontal cortex. We therefore elected to use 4 days as the treatment duration for our studies.

Upon repeated (twice a day for 4 days) administration (s.c.) of corticosterone (50 mg/kg), we found up-regulation of *5-HT_2A_R* mRNA expression in the mouse frontal cortex, an effect that was not observed with the closely related *5-HT_2C_R* or the *dopamine D_2_ receptor* (*D_2_R*) (Fig. 4A). Additionally, this paradigm of repeated corticosterone administration led to a decrease of GR enrichment at its predicted binding site within the *5-HT_2A_R* promoter (Fig. 4B), an effect that was not observed at a region within exon 1 (Fig. 4C).

**Fig. 4.**
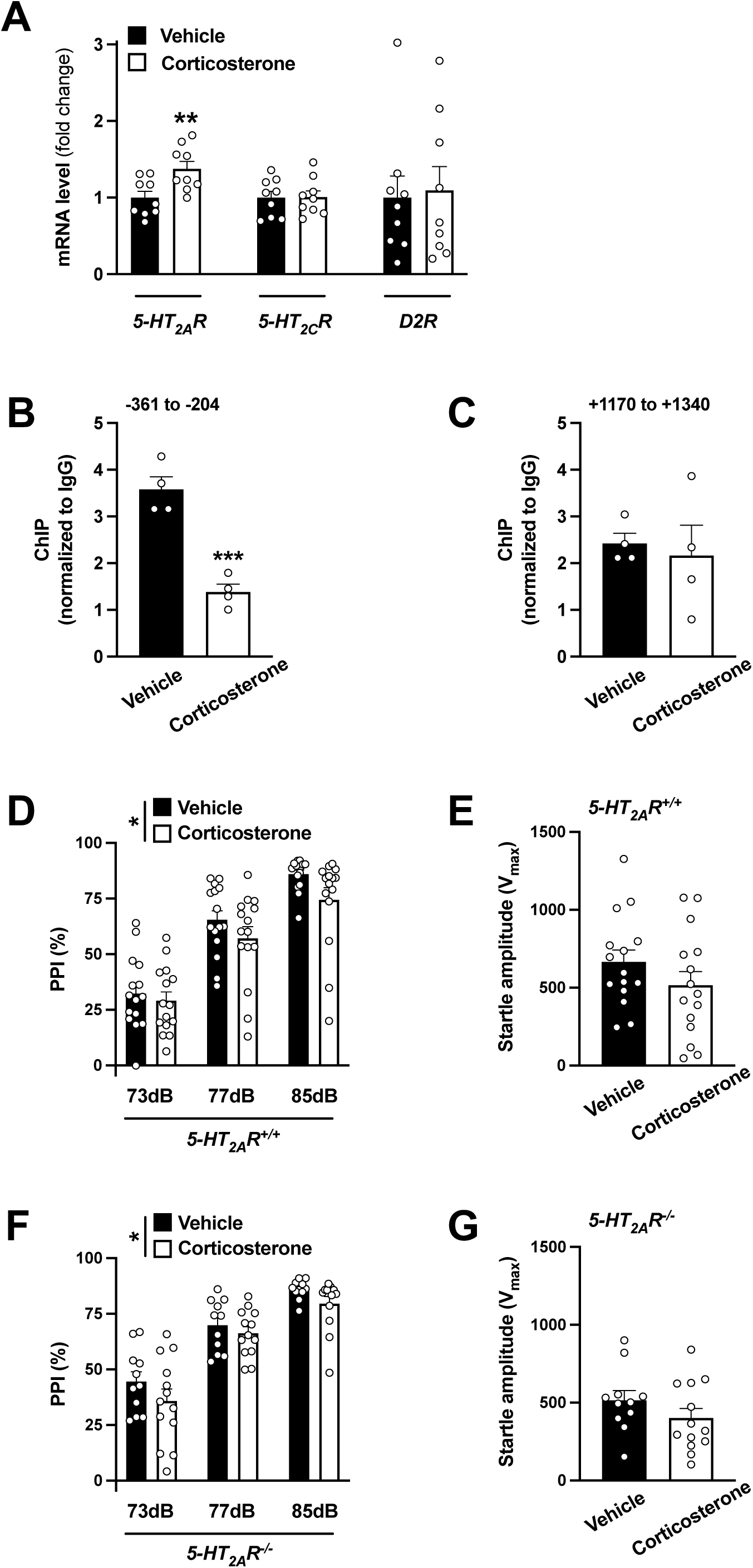
Repeated corticosterone administration up-regulates frontal cortex *5-HT_2A_R* mRNA expression and disrupts sensorimotor gating in mice. **A**, Increased expression of *5-HT_2A_R* mRNA in frontal cortex after repeated corticosterone administration. Mice were treated (twice a day for 4 days) with corticosterone (50 mg/kg; s.c.) or vehicle, and sacrificed for analysis 8.5-13.5 h after the last injection. Expression of *5-HT_2A_R*, *5-HT_2C_R* and *D_2_R* mRNAs in frontal cortex was assessed by qRT-PCR (n = 9 male 129S6/SvEv mice/group). **B-C**, Enrichment of the GR at the region of the *5-HT_2A_R* gene containing a putative GR binding site (**B**), but not within exon 1 (**C**), is decreased upon repeated corticosterone administration in mouse frontal cortex (n = 4 male 129S6/SvEv mice/group). **D-G**, Effect of repeated corticosterone administration on %PPI (**D,F**) and startle amplitude (**E,G**) in *5-HT_2A_R^+/+^* (n = 15 male 129S6/SvEv mice/group) and *5-HT_2A_R^-/-^* (n = 11-13 male 129S6/SvEv mice/group) animals. Data show mean ± S.E.M. *p < 0.05, **p < 0.01. Unpaired two-tailed student’s *t*-test (**A**: *5-HT_2A_R* t_16_ = 2.98, p < 0.01; *5-HT_2C_R* t_16_ = 0.06, p > 0.05; *D_2_R* t_16_ = 0.23, p > 0.05; **B**: t_6_ = 6.95, p < 0.001; **C**: t_6_ = 0.38, p > 0.05; **E**: t_28_ = 1.28, p > 0.05; **G**: t_22_ = 1.30, p > 0.05). Two-way ANOVA followed by Bonferroni’s post-hoc test (**D**: corticosterone vs vehicle F[1, 84] = 4.87, p < 0.05; prepulse intensity F[2, 84] = 67.28, p < 0.001; **F**: corticosterone vs vehicle F[1, 66] = 4.20, p < 0.05; prepulse intensity F[2, 66] = 66.62, p < 0.001; for three-way ANOVA analysis, see Table S4).

Using the same model of repeated corticosterone administration, we next tested for the effects on sensorimotor gating and the role of 5-HT_2A_R-dependent signaling mediating this behavior. Corticosterone treatment reduced %PPI in *5-HT_2A_R^+/+^* mice (Fig. 4D), an effect that was also observed in *5-HT_2A_R^-/-^* littermates (Fig. 4F). Startle magnitude was unaffected upon repeated corticosterone treatment (Figs. 4E,G), whereas *5-HT_2A_R^-/-^* mice showed a statistically significant increase in %PPI as a genotype effect compared to *5-HT_2A_R^+/+^* controls (Figs. 4D,F and Table S4). These data suggest that although repeated corticosterone treatment augments *5-HT_2A_R* mRNA expression in mouse frontal cortex – an effect that occurs in parallel with decreased binding of the GR the *5-HT_2A_R* promoter – the negative effect of this repeated pharmacological activation of the GR on sensorimotor gating processes does not seem to be dependent on 5-HT_2A_R-dependent signaling mechanisms.

Systemically administered glucocorticoids exert a wide variety of physiological effects and pharmacological actions. To further test the functional significance of changes in frontal cortex GR-dependent transcriptional activity on sensorimotor gating processes via 5-HT_2A_R, we next tested the effects of AAV-mediated gene transfer of a previously described GR construct (ΔGR) [67, 68], which lacks the hormone binding domain and is therefore able to constitutively translocate to the nucleus where it has the potential to affect transcription. The ΔGR construct generated from mouse cDNA was subcloned into the AAV8 viral vector under the control of the *CaMKIIα* promoter, which, as mentioned above, allows selective expression of the transgene in frontal cortex glutamatergic pyramidal, but not GABAergic inhibitory, neurons. Additionally, the inclusion of the p2A peptide allows for expression of ΔGR and eYFP as independent proteins from the same transcript [69] (Fig. 5A). This was validated by immunocytochemical (Fig. S6A) and immunoblot assays in Neuro-2a cells (Fig. 5C), which endogenously express the *CaMKIIα* gene, as well as immunohistochemical assays in mouse frontal cortex tissue sections (Fig. 5B). First, we confirmed that mice injected with the AAV-CaMKII*α*:ΔGR-p2A-eYFP construct (AAV-ΔGR-eYFP) showed elevated expression of *GR* in the frontal cortex (Fig. 5D), as well as upregulation of *FKBP5* (Fig. S6C), but not *FKBP4* (Fig. S6B), mRNA expression. Notably, such overexpression of ΔGR decreased *5-HT_2A_R* mRNA, but not *5-HT_2C_R* or *D_2_R* mRNAs in this brain region (Fig. 5E).

**Fig. 5.**
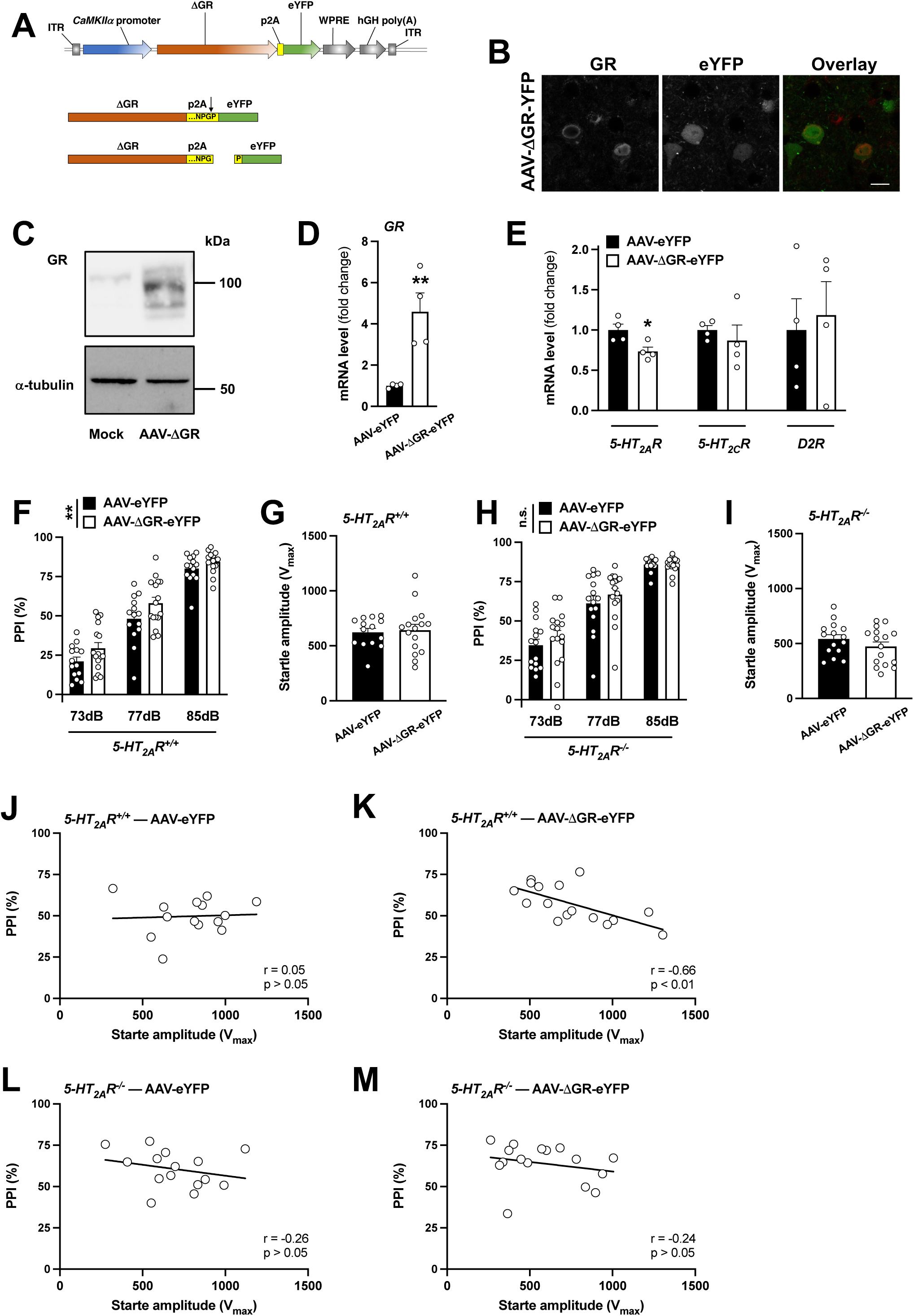
Virally-mediated augmentation of GR function disrupts sensorimotor gating via 5-HT_2A_R. **A,** Schematic representation of the recombinant AAV vector used to over-express both ΔGR and eYFP under the *CaMKIIα* promoter in mouse frontal cortex pyramidal neurons. The arrowhead indicates p2A cleave site. **B,** Representative immunohistochemical images of the mouse frontal cortex injected with the AAV-CaMKII*α*::ΔGR-p2A-eYFP (AAV-ΔGR-eYFP) construct. Note that signals of ΔGR (red) and eYFP (green) do not overlap, indicating high cleavage efficacy of the p2A site. **C,** Representative immunoblot of Neuro-2a cells transfected with mock or AAV-ΔGR-p2A-eYFP plasmids. **D-E,** AAV-ΔGR-eYFP or AAV-eYFP empty vector was injected into the frontal cortex. Mice were sacrificed for analysis 3 weeks after surgery, and expression of *GR* mRNA (**D**) and *5-HT_2A_R*, *5-HT_2C_R* and *D_2_R* mRNAs (**E**) in frontal cortex was assessed by qRT-PCR (n = 4 male and female 129S6/SvEv mice/group). **F-I,** AAV-ΔGR-eYFP or AAV-eYFP empty vector was injected into the frontal cortex of *5-HT_2A_R^+/+^* (n = 14-16 male 129S6/SvEv mice/group) and *5-HT_2A_R^-/-^* (n = 15-16 male 129S6/SvEv mice/group) animals. Behavior was tested 3 weeks after surgery**. J-M,** Effect of AAV-ΔGR-eYFP (**J,L**) vs AAV-eYFP (**K,M**) on Pearson’s correlation between startle magnitudes within the 5 stimulus-only trials at the beginning of the PPI session and %PPI in *5-HT_2A_R^+/+^* (**J,K**) and *5-HT_2A_R^-/-^* (**L,M**) mice. Data show mean ± S.E.M. *p < 0.05, **p < 0.01, n.s., not-significant. Unpaired two-tailed student’s *t*-test (**D**: t_6_ = 3.93, p < 0.01; **E**: *5-HT_2A_R* t_6_ = 2.93, p < 0.05; *5-HT_2C_R* t_6_ = 0.66, p > 0.05; *D_2_R* t_6_ = 0.32, p > 0.05; **G**: t_28_ = 0.27, p > 0.05; **I**: t_29_ = 1.22, p > 0.05). Two-way ANOVA followed by Bonferroni’s post-hoc test (**F:** AAV-ΔGR-eYFP vs AAV-eYFP F[1, 84] = 8.30, p < 0.01; prepulse intensity F[2, 84] = 159.6, p < 0.001; **H:** AAV-ΔGR-eYFP vs AAV-eYFP F[1, 87] = 1.32, p > 0.05; prepulse intensity F[2, 87] = 92.77, p < 0.001; for three-way ANOVA analysis, see Table S5). Correlation analysis was conducted using Pearson’s *r* (**J-M**). Scale bar, 10 µm (**B**).

To determine whether overexpression of ΔGR influences behavioral responses that require 5-HT_2A_R-mediated signaling pathways, we tested the effects of AAV-ΔGR-eYFP compared to the empty AAV-eYFP vector on sensorimotor gating behavior in *5-HT_2A_R^+/+^* and *5-HT_2A_R^-/-^* mice. As before (Figs. 4D,F and Table S4), %PPI was increased in *5-HT_2A_R^-/-^* animals as compared to *5-HT_2A_R^+/+^* controls (Figs. 5F,H and Table S5), whereas startle amplitude was similar in both groups of mice (Figs. 5G,I). Notably, AAV-mediated overexpression of ΔGR in the frontal cortex increased %PPI behavior in *5-HT_2A_R^+/+^* animals (Fig. 5F), whereas a similar manipulation of GR-dependent function failed to affect sensorimotor gating processes when this behavior model was tested in *5-HT_2A_R^-/-^* mice (Fig. 5H).

We also tested whether AAV-dependent manipulation of GR transcriptional activity affected the correlation between the magnitude of the startle amplitudes during the 5 pulse-only trials at the beginning of the PPI session and the average %PPI in both genotypes. Importantly, frontal cortex injection of the AAV-ΔGR-eYFP construct led to a negative correlation between startle amplitudes prior to the PPI test and average %PPI in *5-HT_2A_R^+/+^* mice (Figs. 5J,K), an effect that was not observed in *5-HT_2A_R^-/-^* littermates (Figs. 5L,M). This suggests that virally mediated augmentation of frontal cortex GR function dominates behavioral sensorimotor gating processes via 5-HT_2A_R.

## DISCUSSION

Prior to these studies, although 5-HT_2A_R dysregulation in frontal cortex of rodent MIA models had been demonstrated [28–30], the underlying cause of this alteration remained unsolved. Our data using three independent models suggest that alterations in GR signaling, including both decreased nuclear immunoreactivity and reduced GR binding to the *5-HT_2A_R* promoter, underlie increased *5-HT_2A_R* expression in MIA offspring mouse frontal cortex. This 5-HT_2A_R dysregulation, in turn, was necessary for MIA-induced reduction in mature mushroom frontal cortex dendritic spine density – implicating this process of synaptic structure alterations in the MIA model. Since similar GR alterations were observed in frontal cortex of postmortem schizophrenia samples, these studies provide key insights into mechanisms leading to 5-HT_2A_R dysregulation in the context of prenatal insults, clarifying the GR-dependent control of *5-HT_2A_R* abnormal expression and elucidating potential mechanisms involved in synaptic and sensorimotor gating deficits within the preclinical model.

One of the most intriguing findings of our study was the potential repressive role of the transcription factor GR in the promoter activity of the *5-HT_2A_R* gene. Using ChIP assays in frontal cortex samples from naïve mice, we showed enrichment of GR binding at a region of the *5-HT_2A_R* gene containing a putative GR binding site. However, in frontal cortex samples of adult mice born to mothers injected with poly-(I:C) during pregnancy, we found a significantly decreased enrichment of GR at the *5-HT_2A_R* gene promoter. This MIA-dependent epigenetic modification was associated with augmentation of *5-HT_2A_R* mRNA transcription in the mouse frontal cortex, which suggests that GR plays a repressive role in the transcriptional activity of the *5-HT_2A_R* gene. This concept is further supported by our findings suggesting downregulation of the *5-HT_2A_R* expression upon continued activation of the GR pathway via AAV-ΔGR, which provides a crucial advancement to work on the relationship between the two receptors, and yields information about a potential mechanism for the exquisite sensitivity of 5-HT_2A_Rs to stress.

Our current data raise the question of why repeated administration of systemic corticosterone and local frontal cortex AAV-ΔGR injection exert opposite effects on both *5-HT_2A_R* mRNA expression and %PPI. One possible explanation is the ability of negative feedback to constrain GR-dependent signaling. Although both repeated corticosterone and ΔGR result in increased *FKBP5* expression, suggesting the existence of negative feedback within both experimental systems, AAV-ΔGR injection results in a roughly fourfold, at least at the mRNA level, induction of *GR* expression. In addition, the ΔGR truncation is capable of constitutive nuclear translocation, and is not expressed under the control of the endogenous *GR* locus, both of which could render its resistance to endogenous negative feedback mechanisms. Our observation that repeated corticosterone administration results in decreased enrichment of the GR at the *5-HT_2A_R* promoter further supports the hypothesis that negative feedback may explain this discrepancy. Thus, sustained AAV-mediated ΔGR occupancy at the *5-HT_2A_R* promoter would suppress *5-HT_2A_R* transcription while negative feedback upon repeated corticosterone administration would reduce endogenous GR occupancy of the *5-HT_2A_R* promoter and allow stimulation of *5-HT_2A_R* expression. In addition, while ΔGR-induced PPI improvements were absent in *5-HT_2A_R^-/-^* mice, corticosterone-induced PPI deficits were observable in both genotypes. While this might be explained by the systemic nature of corticosterone administration, in contrast to the frontal cortex-specific AAV-mediated ΔGR expression, further studies are needed to clarify what molecular underpinnings sensorimotor gating alterations induced by this model of repeated pharmacological activation of the GR.

We report that MIA-induced alterations in mushroom dendritic spine density are 5-HT_2A_R-dependent. Of note, *5-HT_2A_R^-/-^* mice exhibited lower mature mushroom spine density compared to *5-HT_2A_R^+/+^* controls regardless of the maternal treatment. Within a different experimental paradigm, we have recently observed similar decreases in frontal cortex spine density in *5-HT_2A_R^-/-^* mice as compared to *5-HT_2A_R^+/+^* controls [44]. Given the implication of ligands that activate [44, 70] or block [43] 5-HT_2A_R-dependent signaling in processes related to synaptic structural and functional plasticity, our findings corroborate the fundamental role of this serotonin receptor in these processes.

Our data support a negative regulatory relationship between GR-dependent signaling and 5-HT_2A_R expression with implications for dendritic structural plasticity, functional GR nuclear translocation and behavior phenotypes observed in mouse MIA models. These findings advance our understanding of the molecular mechanisms by which prenatal insults up-regulate 5-HT_2A_R expression in the frontal cortex of the adult offspring, which may provide the basis for the design of alternative approaches to reduce the impact of MIA and other early life events on physiology and behavior.

## ACKNOWLEDGEMENTS

The authors thank Mario de la Fuente for his critical review of the manuscript, Terrell Holloway for his early help in PPI experiments, and the staff members of the Basque Institute of Legal Medicine for their cooperation in the study.

## AUTHOR CONTRIBUTIONS

J.M.S. and J.G.-M. conceived and designed the experiments, analyzed the data, and wrote the manuscript. J.M.S. performed experiments. C.M., supervised by J.J.M., performed assays in postmortem human brain samples. S.S. helped with FISH assays. J.L.M. performed initial dendritic spine structure assays. L.F.C. and J.J.M. obtained and classified postmortem human brain samples. P.M.B. provided advice on behavioral assays. J.G.-M. supervised the research and obtained funding. All authors discussed the results and commented on the manuscript prior to submission for publication consideration.

---

**Fig. S1.**
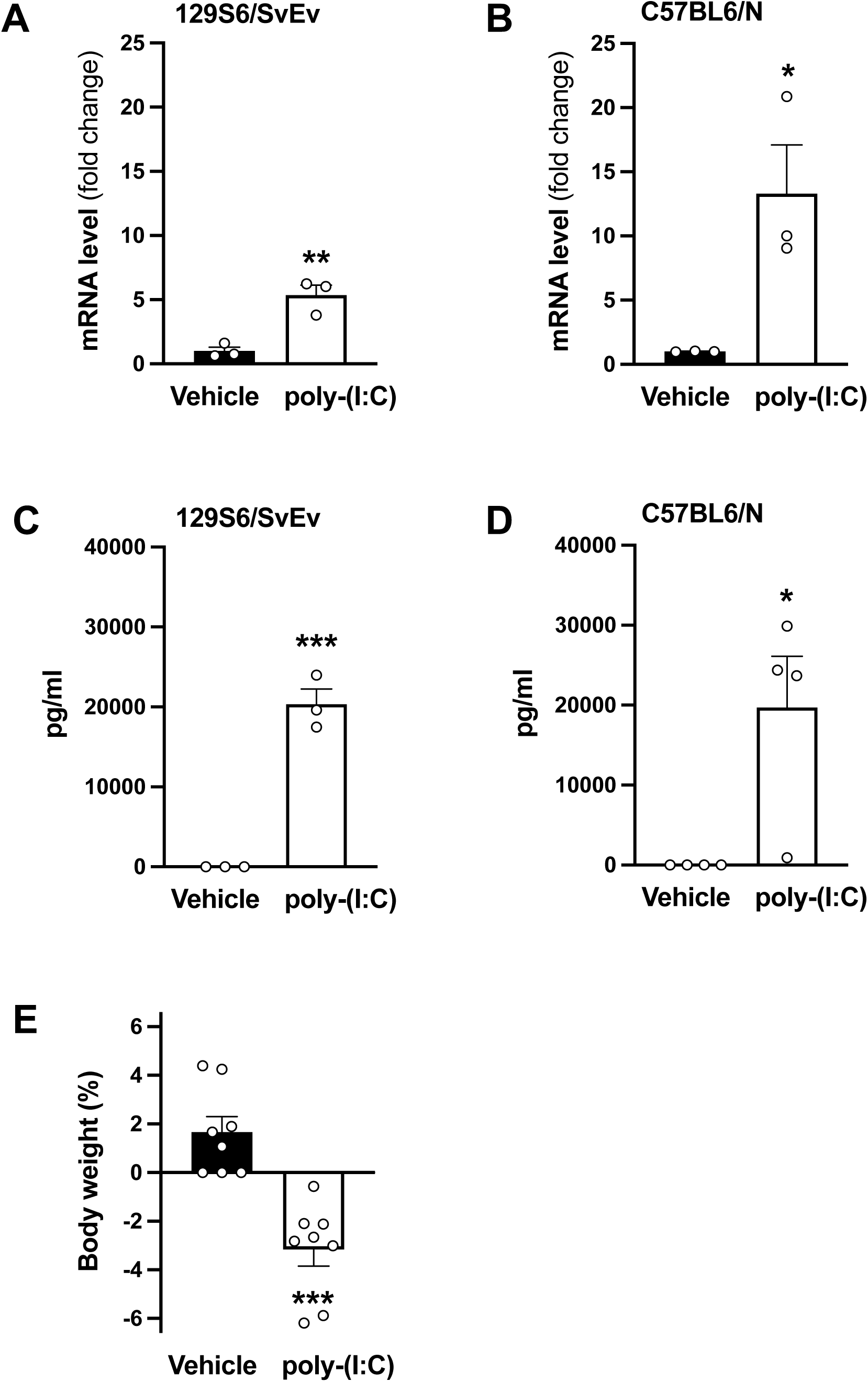
Poly-(I:C) induces an immune response in non-pregnant female mice. **a-d,** Adult female 129S6/SvEv (in-house) (**A,C**) and C57BL6/N (CRL) (**B,D**) mice were injected (i.p.) with poly-(I:C) (20 mg/kg) or vehicle, and sacrificed 2.5 after administration. *IL-6* mRNA was assessed by qRT-PCR in mouse frontal cortex samples (**A,B**), and IL-6 immunoreactivity was assessed by ELISA assays in serum samples (**C,D**) (n = 3-4 female mice/group). **E,** Adult female C57BL6/N (CRL) mice were injected (i.p.) with poly-(I:C) (20 mg/kg) or vehicle, and percentage change in weight vas evaluated 24 h after (n = 8 female mice/group). Data show mean ± S.E.M. *p < 0.05, **p < 0.01, ***p < 0.001. Unpaired two-tailed student’s *t*-test (**A**: t_4_ = 5.21, p < 0.01; **B**: t_4_ = 3.25, p < 0.05; **C:** t_4_ = 10.65, p < 0.001; **D**: t_4_ = 3.07, p < 0.05; **E**: t_4_ = 5.17, p < 0.001).

**Fig. S2.**
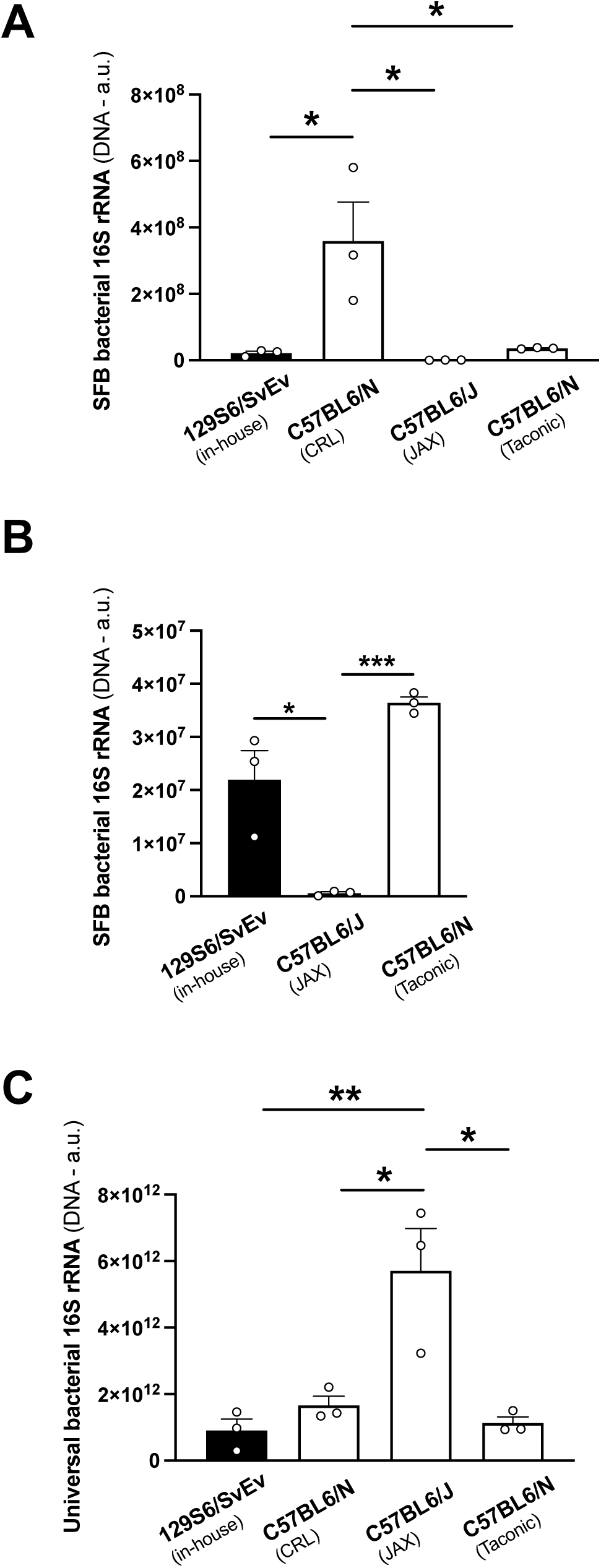
Gut microbiota are enriched for segmented filamentous bacteria (SFB) in 129S6/SvEv (in-house), C57BL6/N (CRL) and C57BL6/N (Tac), but not C57BL6/J (JAX) adult female mice. **A-C**, DNA was extracted from mouse cecum contents and the 16S rRNA gene was assessed by qPCR using primer pairs targeting either SFB (**A**) or universal primer pairs that detect this gene across the vast majority of bacterial taxa (**C**) (n = 3 female mice/group). **B**, The same data as in “A” are depicted, but the C57BL6/N (CRL) mice were excluded from the data analysis. Data show mean ± S.E.M. *p < 0.05, **p < 0.01, ***p < 0.001. Two-way ANOVA followed by Bonferroni’s post-hoc test (**A**: F[3, 8] = 8.40, p < 0.01; **B**: F[2, 6] = 30.95, p < 0.001; **C**: F[3, 8] = 11.9, p < 0.01).

**Fig. S3.**
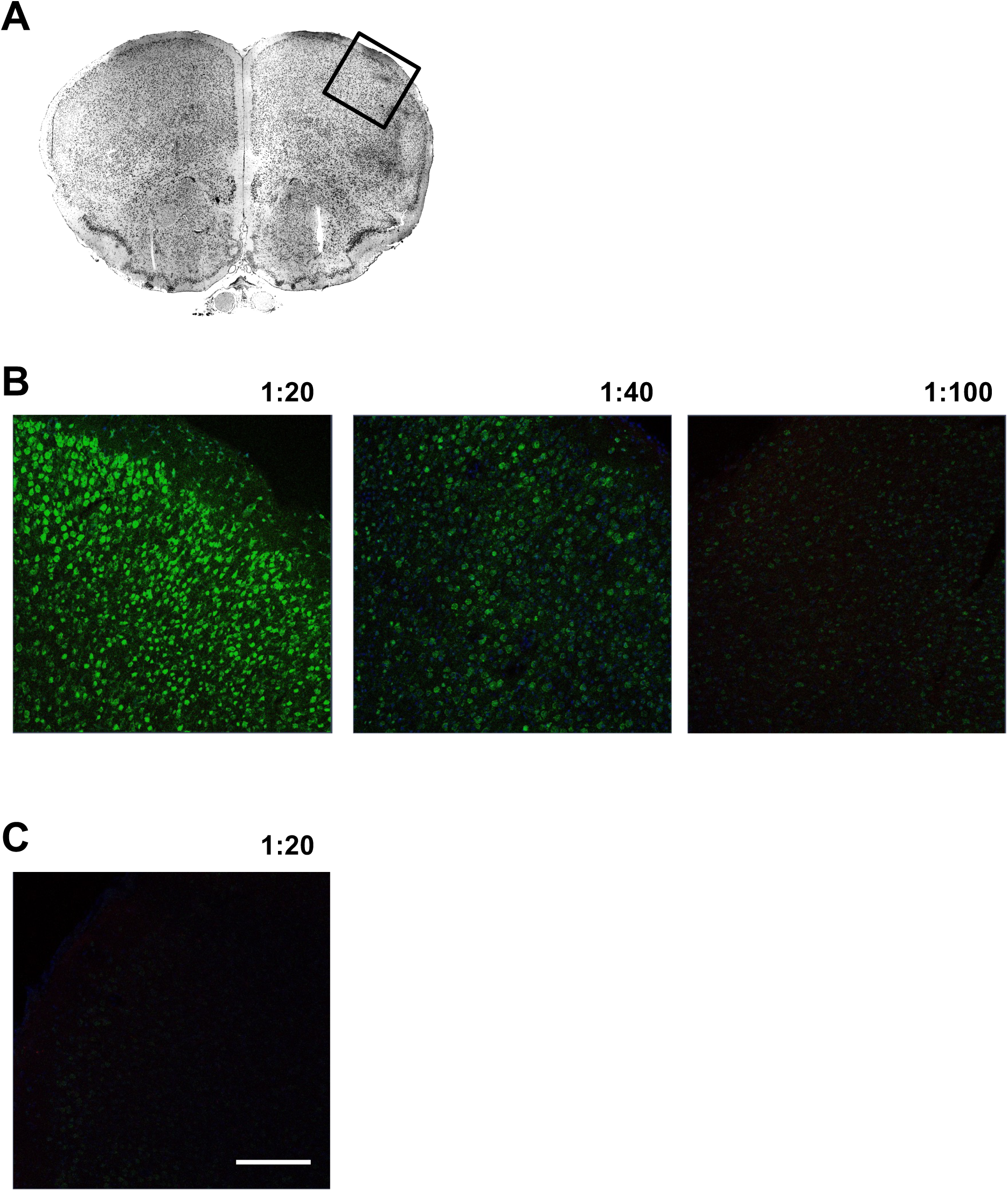
Control assays validate the FISH assay with the antisense *5-HT_2A_R* probe in mouse frontal cortex. **A,** Nissl-staining image of mouse frontal cortex for neuroanatomical orientation. **B,** FISH signal in male C57BL6/N CRL mouse frontal cortex decreases with increasing dilution factors of the antisense *5-HT_2A_R* probe. **C,** Signal is absent with a sense *5-HT_2A_R* probe. Scale bar, 200 µm (**A**).

**Fig. S4.**
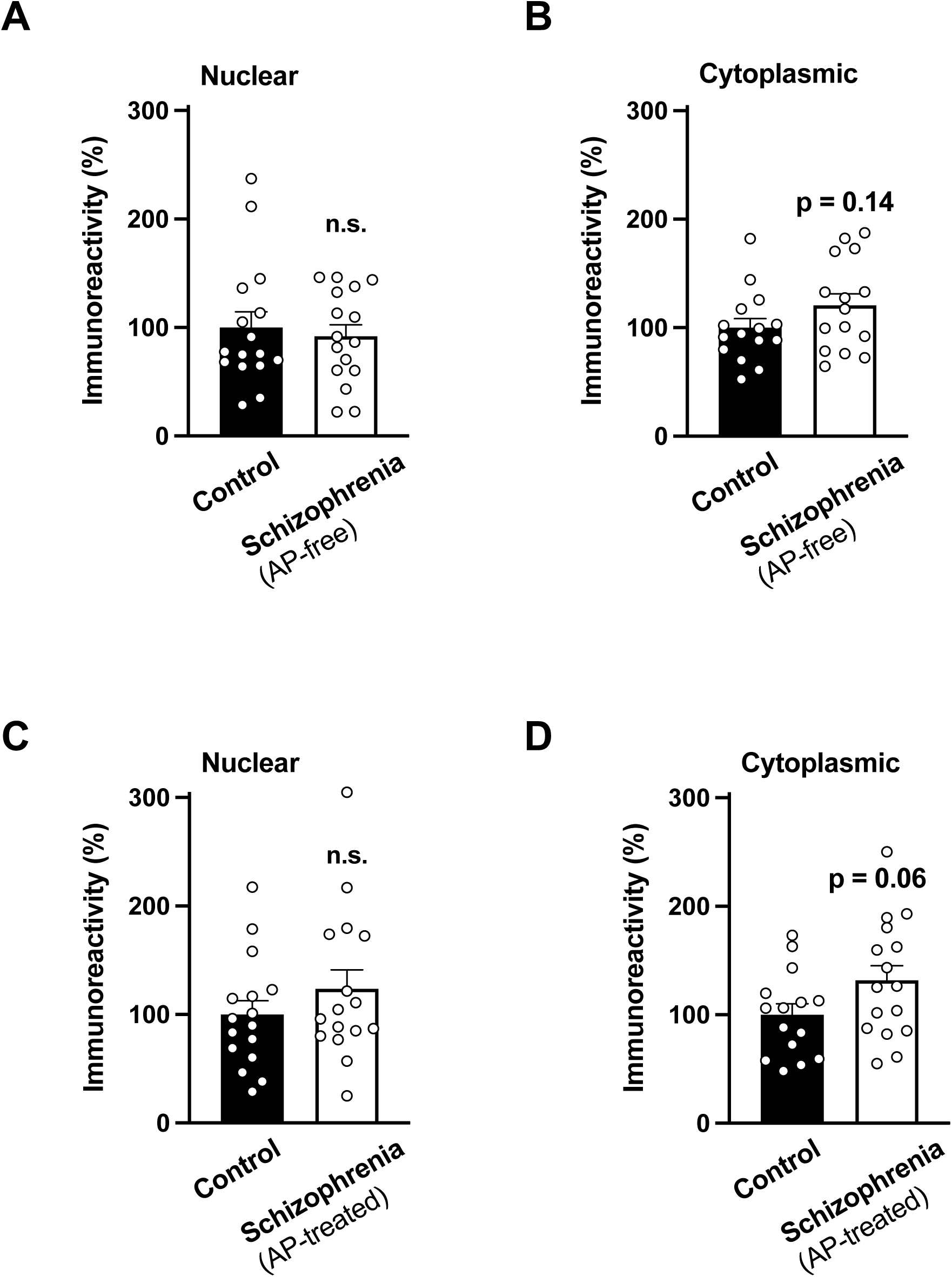
GR localization is dysregulated in postmortem frontal cortex of schizophrenia subjects. **A-D,** Immunoblots showed a trend towards upregulation of GR protein in cytoplasmic (**B,D**) but not nuclear (**A,C**) preparations from postmortem human brain samples of antipsychotic-free (AP-free) (**A,B**) and antipsychotic-treated (AP-treated) (**C,D**) schizophrenia subjects as compared to controls (for demographic information, see Tables S2 and S3). Data show mean ± S.E.M. n.s., not significant. Unpaired two-tailed student’s *t*-test (**A:** t_30_ = 0.45, p > 0.05; **B**: t_28_ = 1.49, p = 0.14; **C:** t_30_ = 1.09, p > 0.05; **D:** t_29_ = 1.85, p = 0.06).

**Fig. S5.**
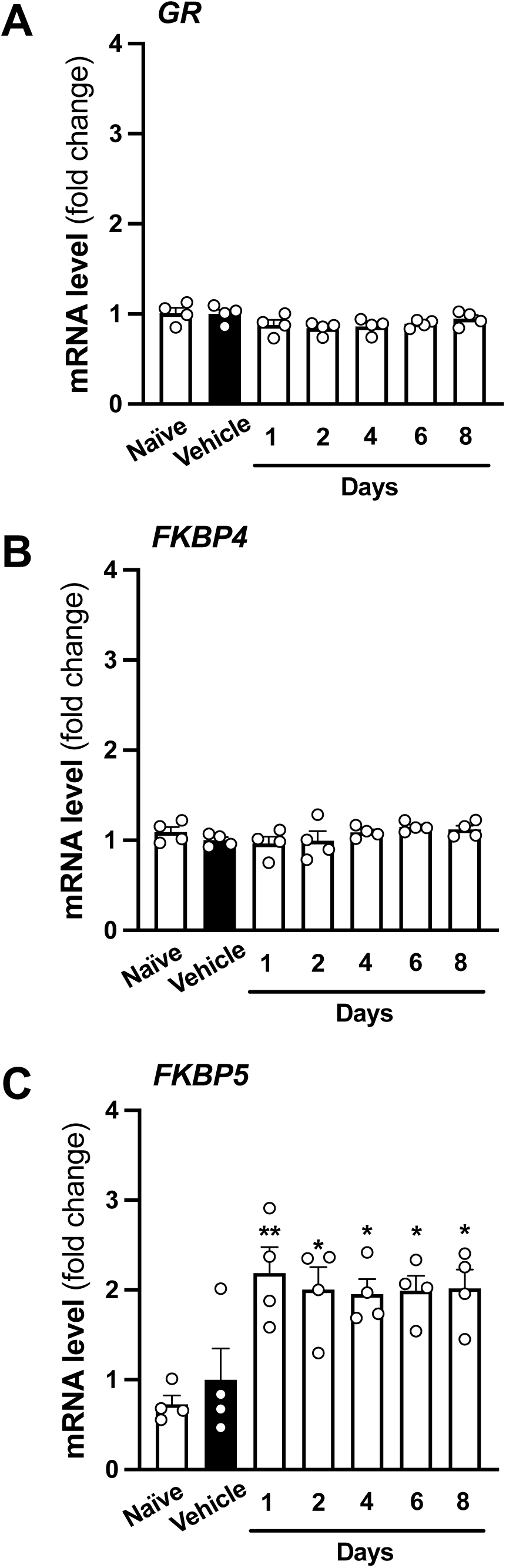
A time-course of repeated corticosterone administration reveals *FKBP5* mRNA induction in mice. **A-C,** Adult male mice were administered (s.c.) corticosterone (50 mg/kg) twice a day for either 1, 2, 4, 6 or 8 days, and sacrificed to collect samples 8.5-13 h after the last injection. Control groups included mice injected with vehicle as well as injection-naïve mice. Expression of *GR* (**A**) *FKBP4* (**B**), and *FKBP5* (**C**) mRNAs in frontal cortex was assessed by qRT-PCR (n = 4 male 129S6/SvEv in-house mice/group). Data show mean ± S.E.M. *p < 0.05, **p < 0.01. Two-way ANOVA followed by Bonferroni’s post-hoc test (**A:** F[6, 21] = 2.17, p = 0.087; **B:** F[6, 21] = 1.38, p > 0.05; **C:** F[6.21] = 6.26, p < 0.001).

**Fig. S6.**
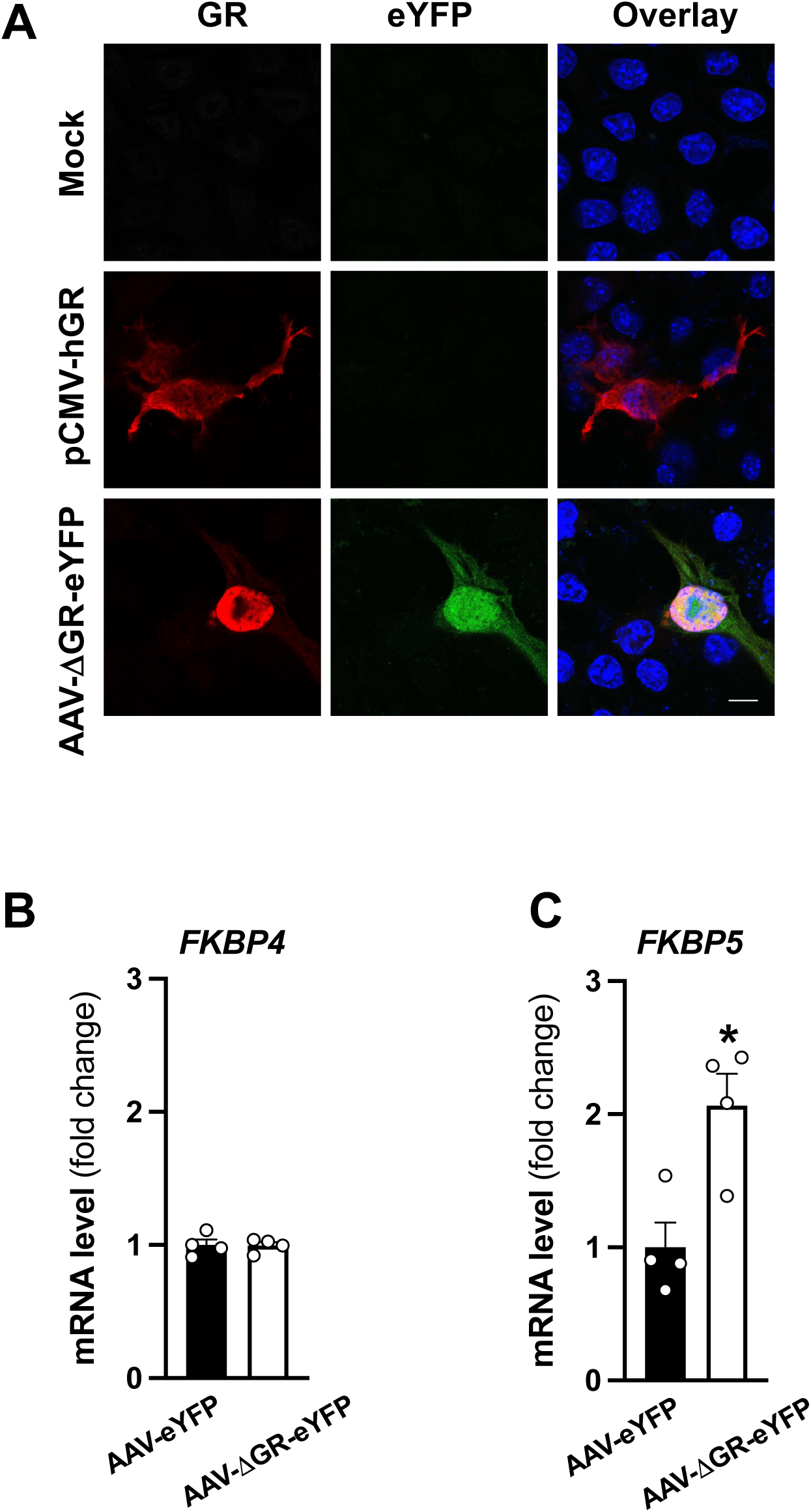
The ΔGR construct is expressed in Neuro-2a cells and in mouse frontal cortex. **A,** Representative immunocytochemical images of Neuro-2a cells transfected with mock, pCMV-hGR or AAV-ΔGR-p2A-eYFP plasmids. **B-C,** AAV-ΔGR-eYFP or AAV-eYFP empty vector was injected into the frontal cortex. Mice were sacrificed for analysis 3 weeks after surgery, and expression of *FKBP4* (**B**) and *FKBP5* (**C**) in frontal cortex was assessed by qRT-PCR (n = 4 male 129S6/SvEv mice/group). Data show mean ± S.E.M. *p < 0.05. Unpaired two-tailed student’s *t*-test (**B**: t_6_ = 0.098, p > 0.05; **C**: t_6_ = 3.51, p < 0.05). Nuclei were stained in blue with Hoechst (**A**). Scale bar, 10 µm (**A**).

**Table S1.**
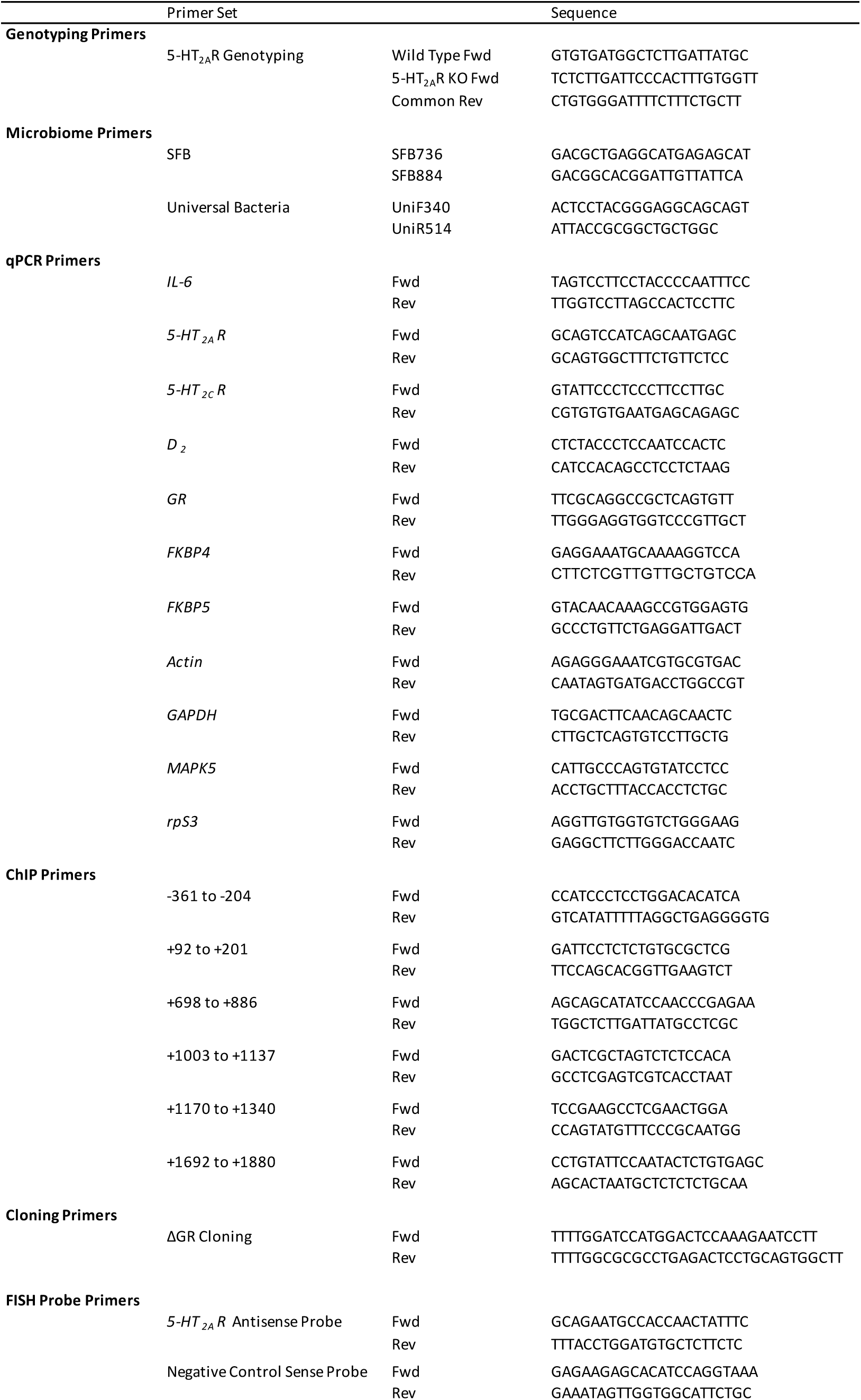
List of primer pairs

**Table S2.** Demographic characteristics of antipsychotic-free schizophrenia subjects and their respective control subjects.

|  | **Sex**<br>**(F/M)** | **Age at**<br>**death**<br>**(years)** | **Postmorte**<br>**m delay (h)** | **Storage**<br>**time**<br>**(months)** | **Antipsychoti**<br>**c treatment** | **Additional**<br>**drugs** |
| --- | --- | --- | --- | --- | --- | --- |
| Schizophrenia 1 | M | 30 | 51 | 203 | (-) | (-) |
| Control 1 | M | 29 | 18 | 68 |  | (-) |
| Schizophrenia 2 | M | 48 | 20 | 122 | (-) | (-) |
| Control 2 | M | 47 | 18 | 142 |  | BDZ |
| Schizophrenia 3 | M | 31 | 11 | 113 | (-) | (-) |
| Control 3 | M | 31 | 13 | 99 |  | ETH (0.96 g/l) |
| Schizophrenia 4 | M | 45 | 3 | 106 | (-) | BDZ |
| Control 4 | M | 44 | 21 | 92 |  | (-) |
| Schizophrenia 5 | M | 27 | 24 | 104 | (-) | (-) |
| Control 5 | M | 28 | 30 | 178 |  | (-) |
| Schizophrenia 6 | M | 46 | 22 | 69 | (-) | (-) |
| Control 6 | M | 46 | 24 | 67 |  | (-) |
| Schizophrenia 7 | M | 48 | 11 | 50 | (-) | (-) |
| Control 7 | M | 49 | 8 | 43 |  | (-) |
| Schizophrenia 8 | M | 45 | 18 | 59 | (-) | (-) |
| Control 8 | M | 47 | 15 | 44 |  | (-) |
| Schizophrenia 9 | M | 34 | 15 | 62 | (-) | (-) |
| Control 9 | M | 36 | 48 | 117 |  | (-) |
| Schizophrenia 10 | M | 52 | 7 | 66 | (-) | BDZ |
| Control 10 | M | 51 | 13 | 46 |  | (-) |
| Schizophrenia 11 | M | 49 | 12 | 33 | (-) | BDZ |
| Control 11 | M | 46 | 6 | 28 |  | ETH (1.91 g/l) |
| Schizophrenia 12 | M | 60 | 12 | 43 | (-) | BDZ |
| Control 12 | M | 62 | 9 | 66 |  | (-) |
| Schizophrenia 13 | F | 67 | 22 | 52 | (-) | (-) |
| Control 13 | F | 66 | 16 | 24 |  | (-) |
| Schizophrenia 14 | M | 34 | 23 | 116 | (-) | (-) |
| Control 14 | M | 33 | 28 | 28 |  | ETH (0.97 g/l) |
| Schizophrenia 15 | M | 32 | 52 | 121 | (-) | BDZ |
| Control 15 | M | 32 | 4 | 44 |  | ETH (2.98 g/l) |
| Schizophrenia 16 | M | 32 | 21 | 122 | (-) | (-) |
| Control 16 | M | 32 | 16 | 33 |  | (-) |
| Schizophrenia | 1F/15<br>M | 42.5±2.9 | 20.3 ±3.4 | 90.1±10.9 |  |  |
| Control | 1F/15<br>M | 42.4±2.9 | 17.9±2.7 | 69.9±11.2 |  |  |

Antipsychotics were not detected in blood samples of schizophrenia subjects at the time of death. Abbreviations: benzodiazepines (BDZ), ethanol (ETH).

**Table S3.** Demographic characteristics of antipsychotic-treated schizophrenia subjects and their respective control subjects.

|  | **Sex**<br>(F/M) | **Age at death**<br>(years) | **Postmortem delay (h)** | **Storage time**<br>(months) | **Antipsychotic treatment** | **Additional drugs** |
| --- | --- | --- | --- | --- | --- | --- |
| Schizophrenia 17 | M | 30 | 18 | 130 | OLZ | (-) |
| Control 17 | M | 29 | 36 | 185 |  | ETH (1.71 g/l) |
| Schizophrenia 18 | M | 32 | 8 | 125 | QTP | BDZ |
| Control 18 | M | 32 | 4 | 24 |  | (-) |
| Schizophrenia 19 | M | 23 | 16 | 120 | SUL | (-) |
| Control 19 | M | 23 | 17 | 93 |  | (-) |
| Schizophrenia 20 | M | 35 | 3 | 120 | QTP | BDZ |
| Control 20 | M | 36 | 23 | 63 |  | (-) |
| Schizophrenia 21 | M | 33 | 23 | 107 | CLZ | (-) |
| Control 21 | M | 33 | 4 | 99 |  | (-) |
| Schizophrenia 22 | M | 35 | 11 | 70 | CLZ | BDZ |
| Control 22 | M | 36 | 18 | 140 |  | ETH (1.69 g/l) |
| Schizophrenia 23 | F | 42 | 38 | 129 | CLZ, QTP, SUL | BDZ |
| Control 23 | F | 38 | 22 | 46 |  | (-) |
| Schizophrenia 24 | M | 37 | 8 | 75 | OLZ | BDZ, ETH (0.90 g/l) |
| Control 24 | M | 41 | 24 | 180 |  | (-) |
| Schizophrenia 25 | F | 60 | 23 | 54 | CLZ, SUL | BDZ |
| Control 25 | F | 60 | 48 | 60 |  | BDZ |
| Schizophrenia 26 | F | 56 | 13 | 119 | CLZ | (-) |
| Control 26 | F | 58 | 20 | 176 |  | (-) |
| Schizophrenia 27 | M | 43 | 5 | 21 | OLZ | BDZ |
| Control 27 | M | 44 | 29 | 24 |  | (-) |
| Schizophrenia 28 | M | 58 | 6 | 21 | CLP | (-) |
| Control 28 | M | 57 | 3 | 39 |  | (-) |
| Schizophrenia 29 | M | 51 | 18 | 22 | PAL | BDZ |
| Control 29 | M | 50 | 2 | 33 |  | (-) |
| Schizophrenia 30 | M | 51 | 28 | 25 | OLZ,CLP | BDZ |
| Control 30 | M | 50 | 24 | 20 |  | (-) |
| Schizophrenia 31 | M | 34 | 21 | 28 | HAL | BDZ |
| Control 31 | M | 33 | 17 | 33 |  | (-) |
| Schizophrenia 32 | M | 58 | 16 | 33 | APZ | (-) |
| Control 32 | M | 56 | 15 | 18 |  | (-) |
| Schizophrenia | 3F/13M | 42.4±2.9 | 15.9 ±2.3 | 74.9±11.4 |  |  |
| Control | 3F/13M | 42.3±2.9 | 19.1±3.1 | 77.1±15.3 |  |  |

Therapeutic levels of aripiprazole (APZ), clotiapine (CLP), clozapine (CLZ), haloperidol (HAL), olanzapine (OLZ), paliperidone (PAL), quetiapine (QTP) and sulpiride (SUL) were detected in blood samples of schizophrenia subjects at the time of death. Abbreviations: benzodiazepines (BDZ), ethanol (ETH).

**Table S4.** Three-way ANOVA analysis of the effect of repeated corticosterone vs vehicle treatment on %PPI in male 129S6/SvEv (in-house) mice.

| Source of variation | ANOVA | p value |
| --- | --- | --- |
| %PPI | F[2,150] = 125.7 | p < 0.001 |
| Treatment | F[1,150] = 8.55 | p < 0.01 |
| Genotype | F[1,150] = 6.84 | p < 0.01 |
| %PPI × Treatment | F[2,150] = 0.17 | p > 0.05 |
| %PPI × Genotype | F[2,150] = 0.70, | p > 0.05 |
| Genotype × Treatment | F[1,150] = 0.095 | p > 0.05 |
| %PPI × Genotype × Treatment | F[2,150] = 0.54 | p > 0.05 |

**Table S5.** Three-way ANOVA analysis of the effect of AAV-ΔGR-eYFP vs AAV-eYFP on %PPI in male 129S6/SvEv (in-house) mice.

| Source of variation | ANOVA | p value |
| --- | --- | --- |
| %PPI | F[2,171] = 241.5 | p < 0.001 |
| Viral vector | F[1,150] = 8.55 | p < 0.01 |
| Genotype | F[1,171] = 7.64 | p < 0.001 |
| %PPI × viral vector | F[2,171] = 0.78 | p > 0.05 |
| %PPI × Genotype | F[2,171] = 1.71, | p > 0.05 |
| Genotype × viral vector | F[1,171] = 1.10 | p > 0.05 |
| %PPI × Genotype × viral vector | F[2,171] = 0.0028 | p > 0.05 |

